# Drug self-administration in head-restrained mice for simultaneous multiphoton imaging

**DOI:** 10.1101/2020.10.25.354209

**Authors:** Kelsey M. Vollmer, Elizabeth M. Doncheck, Roger I. Grant, Kion T. Winston, Elizaveta V. Romanova, Christopher W. Bowen, Preston N. Siegler, Ana-Clara Bobadilla, Ivan Trujillo-Pisanty, Peter W. Kalivas, James M. Otis

**Affiliations:** Department of Neuroscience, Medical University of South Carolina, Charleston, SC 29425, USA; School of Pharmacy, University of Wyoming, Laramie, WY 82071, USA; Langara College, Vancouver, British Columbia V5Y 2Z6, Canada; Hollings Cancer Center, Medical University of South Carolina, Charleston, SC 29425, USA

## Abstract

Multiphoton microscopy is one of several new technologies providing unprecedented insight into the activity dynamics and function of neural circuits. Unfortunately, many of these technologies require experimentation in head-restrained animals, greatly limiting the behavioral repertoire that can be studied with each approach. This issue is especially evident in drug addiction research, as no laboratories have coupled multiphoton microscopy with simultaneous intravenous drug self-administration, the gold standard of behavioral paradigms for investigating the neural mechanisms of drug addiction. Such experiments would be transformative for addiction research as one could measure or perturb an array of behavior and drug-related adaptations in precisely defined neural circuit elements over time, including but not limited to dendritic spine plasticity, neurotransmitter release, and neuronal activity. Here, we describe a new experimental assay wherein mice self-administer drugs of abuse while head-restrained, allowing for simultaneous multiphoton imaging. We demonstrate that this approach enables longitudinal tracking of activity in single neurons from the onset of drug use to relapse. The assay can be easily replicated by interested labs for relatively little cost with readily available materials and can provide unprecedented insight into the neural underpinnings of substance use disorder.

## INTRODUCTION

Multiphoton microscopy provides unprecedented insight into the neurobiological substrates that coordinate behavior. By harnessing fluorescent imaging strategies based on genetically encoded constructs, its never-before-seen spatiotemporal resolution in living tissues enables observation and tracking of cellular activity (e.g., calcium ion concentration or voltage changes; Chen et al., 2013; Lin and Schnitzer, 2016; Tian et al., 2009; Villette et al., 2019), neurotransmitter release (e.g., receptor-activation-based neurotransmitter sensors; Feng et al., 2019; Sun et al., 2018) and morphology changes (e.g., dendritic spine plasticity; Moda-Sava et al., 2019; Muñoz-Cuevas et al., 2013) in awake, behaving animals. Additionally, multiphoton microscopy allows optogenetic or lesion-dependent manipulation of neural circuits from the level of defined neuronal ensembles (Carrillo-Reid et al., 2016; Hill et al., 2017) to axons and dendritic spines (Allegra Mascaro et al., 2010; Canty et al., 2013; Park et al., 2019). This combination of tools has been transformative for neuroscience, as scientists can now observe and interrogate adaptations in neural circuits with astounding precision to better understand the nervous system and the neurobiological underpinnings of disease. Multiphoton microscopy could be particularly transformative for studying substance use disorder (SUD), which is rooted in maladaptive neural plasticity in circuits that govern motivated behavior (Russo and Nestler, 2013). Unfortunately, experiments involving multiphoton microscopy generally require head immobilization, which has prevented the integration of multiphoton imaging with preclinical models of SUD.

Perhaps the most powerful preclinical model for SUD involves training animals to voluntarily self-administer drugs of abuse, an approach that has both predictive and construct validity for treatment outcomes and abuse liability (Epstein et al., 2006; Haney and Spealman, 2008; Shalev et al., 2002). In drug self-administration, animals are reinforced for performing an operant task (e.g., lever pressing) by drug delivery (e.g., intravenous), after which seeking behavior is extinguished through reward omission. Animals can then be tested for drug-seeking reinstatement in response to stimuli known to produce craving and relapse in humans (e.g., drug-associated cues, a single dose of the drug, or stressors). Each phase of self-administration can be used to study the cycle of intoxication, withdrawal, and drug seeking, which characterizes SUD (Koob and Le Moal, 1997). Paralleling the transition from first substance use to chronic misuse, unique neural plasticity emerges during each phase of self-administration (Gardner, 2000). By pairing self-administration with bench approaches, critical factors underlying drug-seeking behavior have been identified at the cellular, molecular, and circuit levels (Deadwyler, 2010; Kalivas and Volkow, 2005; Koob and Le Moal, 1997; Russo et al., 2010). Despite these advances, our ability to precisely observe, longitudinally track, and manipulate these implicated factors in live, behaving animals has been limited.

Here we design and develop a model of drug self-administration in head-restrained mice, an assay allows simultaneous multiphoton imaging. Using otherwise similar experimental parameters, we use this new procedure to replicate patterns of psychostimulant (cocaine) and opioid (heroin) seeking and taking that are typically observed in freely moving animals. Specifically, we find that this procedure reproduces patterns of drug self-administration, extinction, and cue-, drug-, and stress-induced reinstatement as observed in freely moving approaches. Furthermore, we provide example data to reveal the feasibility of simultaneous, longitudinal tracking of activity in single neurons throughout all phases of the task. The experimental assay can be replicated by interested laboratories for relatively little cost using readily available materials.

## METHODS

### Subjects

Male and female C57BL/6J mice (8 wks old/20 g minimum at study onset; Jackson Labs) were group housed pre-operatively and single housed post-operatively under a reversed 12:12-hour light cycle (lights off at 8:00am) with access to standard chow and water *ad libitum*. Experiments were performed in the dark phase and in accordance with the NIH Guide for the Care and Use of Laboratory Animals with approval from the Institutional Animal Care and Use Committee at the Medical University of South Carolina.

### Surgeries

#### Head ring implantation

Mice were anesthetized with isoflurane (0.8-1.5% in oxygen; 1L/minute) and placed within a stereotactic frame (Kopf Instruments) for head ring implantation surgeries. Ophthalmic ointment (Akorn), topical anesthetic (2% Lidocaine; Akorn), analgesic (Ketorlac, 2 mg/kg, ip), and subcutaneous sterile saline (0.9% NaCl in water) treatments were given pre- and intra-operatively for health and pain management. A custom-made ring (stainless steel; 5 mm ID, 11 mm OD) was adhered to the skull using dental cement and skull screws. Head rings were scored on the base using a drill for improved adherence. For mice that also underwent multiphoton imaging, during head ring implantation we microinjected a virus encoding the calcium indicator GCaMP6s (AAVdj-CaMK2α-GCaMP6s; UNC Vector Core) into the dorsal medial prefrontal cortex (dmPFC; AP, +1.85mm; ML, −0.50 mm; DV −2.45 mm). Next, a microendoscopic GRIN lens (gradient refractive index lens; 4mm long, 1mm diameter; Inscopix) was implanted dorsal to dmPFC (AP, +1.85mm; ML,−0.50mm; DV,−2.15mm) as previously described (Otis et al., 2017; Resendez et al., 2016). Head rings were scored on the base using a drill for improved adherence. Animals were subjected to intravenous catheterization surgeries either immediately or four weeks after head ring implantation. Following surgeries, mice received antibiotics (Cefazolin, 200 mg/kg, sc).

#### Intravenous catheterization

Mice were anesthetized as above and implanted with indwelling intravenous catheters for drug self-administration. Catheters (Access Technologies #071709H) were custom-designed for mice, with 4.5 cm of polyurethane tubing (0.012 ID/0.025 OD, rounded tip) between the sub-pedestal/indwelling end of the back-mounted catheter port (Plastics One #8I313000BM01) and silicone vessel suture retention bead, from which 1.0 cm extended intravenously via the right external jugular vein toward the right atrium of the heart. Larger tubing (1.5 cm of 0.025 ID/0.047 OD) encased the smaller attached to the pedestal, and components were adhered by ultraviolet curation (catheter construction was based on Doncheck et al., 2018). Catheters were implanted subcutaneously using a dorsal approach, with back mounts externalized through ≤5 mm midsagittal incisions posterior to the scapulae and tubing running from the sub-pedestal base over the right clavicle into the <1 mm incision in the right external jugular vein. Silk sutures adhered the catheter tubing retention bead to the external jugular and negative pressure-induced blood backflow was assured prior to closing skin incisions with nonabsorbable monofilament sutures. Catheters were flushed daily with heparinized saline (60 units/mL, 0.02 mL) to maintain patency. All animals received analgesic, ophthalmic, and antibiotic treatments as described above in addition to topical antibiotic ointment and lidocaine (2%) jelly application to incisions. Mice were allowed to recover for a minimum of one week before behavioral experiments.

### Self-administration chambers

Drug self-administration chambers were custom designed and created using readily available and cost-effective components. Each chamber was a two-door cabinet shelf (24” × 16” × 30”; NewAge Products; #50002) equipped with 1) soundproofing, 2) breadboards, 3) head-fixation station, 4) restraint tube, 5) operant levers, 5) speakers, 6) infusion pump, 7) electronics, and 8) laptop with MATLAB and Arduino software.

1. <u>Soundproofing</u>: Acoustic foam (2.5” × 24” × 18” UL 94, Professional Acoustics; 12” × 12” × 1.5” egg crate acoustic foam tiles, OBCO) was glued to the inner chamber.
2. <u>Breadboards</u>: An aluminum breadboard (12” × 18’ × 1/2") was used as the base to allow for chamber components to be screwed in place (ThorLabs; MB1218). A smaller aluminum breadboard (4” × 6” × 1/2”) was used to secure the head-fixation station into place and to provide a behavioral platform (ThorLabs; MB4).
3. <u>Head-fixation station</u>: Head-fixation stations were custom-made (Clemson University Engineering Department) and attached to the breadboard. Stainless steel inverted square-edged U-frames with slits in the central crossbar allowed for mouse insertion by head rings. A second crossbar clamped the head rings down to prevent head movement.
4. <u>Restraint tube</u>: Partial body restraint was used to avoid self-injury. Slits were drilled into 50 mL conical tubes (Fisher; 01-812-55) to allow for catheter port externalization while mice were partially restrained within head-fixation stations. The open end of the conical faced the lever box and was positioned to allow free range of movement for the front limbs.
5. <u>Operant levers</u>: Two operant levers (Honeywell; 311SM703-T) were cemented into a hollowed-out 12-well plate (Fisher; FB012928) and wired for active/inactive response functionality through the Arduino board. The levers extended outward by 5 cm from this ‘lever box’ and the ends aligned centrally 3.5 cm from the edge of the restraint conical. Animals could reach the levers by extending their forelimbs.
6. <u>Speaker</u>: Arduino-controlled piezo buzzers (Adafruit; #1739) were positioned above the head-fixation frames for conditioned stimulus (CS) audio tones.
7. <u>Infusion pump</u>: Arduino-controlled infusion pumps (Med Associates #PHM-100VS-2) were used for drug delivery.
8. <u>Arduino board</u>: Arduinos (Arduino Uno Rev 3; A000066) interfaced with two electronic breadboards (Debaser Electronics; DE400BB1-1) for control of self-administration equipment.
9. <u>Laptop</u>: A laptop (Lenovo Ideapad 330S; 81F5006GUS) interfaced with other electronics via Arduino. Arduino- (Arduino 1.8.12) and MATLAB-(MathWorks) software were used to control equipment and record behavioral events.

### Behavioral procedure

Mice underwent 3 days of habituation, during which they were head-restrained for 30 minutes in the operant chambers without access to the levers. During acquisition, the operant levers were placed in front of the mice. Pressing the active lever resulted in immediate tone CS presentation (8 kHz, 2 s), followed by a gap in time (trace interval, 1 s), and finally intravenous drug infusion (2 s; 12.5 μl). The trace interval was included to allow isolated detection of sensory cue- and drug infusion-related neuronal activity patterns, as described previously (Otis et al., 2017). Reinforced active lever presses also resulted in a timeout period wherein further active lever presses were recorded but not reinforced with the CS or drug infusion. Inactive lever presses resulted in neither CS nor drug delivery.

#### Heroin self-administration

Mice underwent 14 days of heroin self-administration on a fixed ratio 1 (FR1) schedule of reinforcement using a decreasing dose design (Day 1-2: 0.1 mg/kg/12.5 μl heroin, 10 infusion maximum; Day 3-4: 0.05 mg/kg/12.5 μl heroin, 20 infusion maximum; Day 5-14: 0.025 mg/kg/12.5 μl heroin, 40 infusion maximum), for a maximum of 1 mg/kg of heroin per session (similar to previously described experiments in freely moving mice; Corre et al., 2018). Session durations were ≤ 2 hrs, depending on whether infusion caps were met. Following acquisition, heroin self-administration mice underwent extinction training, wherein active lever presses resulted in neither CS nor drug delivery until extinction criteria were reached. Extinction criteria were determined a priori, defined as (1) ≥ 10 days of extinction training and (2) 2 of the last 3 days resulting in ≤ 20% of the average active lever pressing observed during the last 2 days of acquisition. Mice that did not reach extinction criteria were excluded from analyses (n=4). Following establishment of extinction criteria, mice underwent reinstatement testing using a counterbalanced design, with cue-, drug-, yohimbine-, predator odor-, or saline-induced reinstatement tests administered in a pseudorandom order. For cue-induced reinstatement testing, active lever presses resulted in CS presentation in a manner identical to acquisition, but drug infusions were excluded. For drug-primed reinstatement, heroin (1 mg/kg, i.p.) was delivered immediately prior to the reinstatement session. A heroin-primed reinstatement dose-response pilot study revealed this dose produced responding most comparable to that observed during both acquisition and other reinstatement tests (data not shown). For reinstatement testing in response to the pharmacological stressor yohimbine, yohimbine (0.0625 mg/kg, i.p., 30 min pretreatment; Sigma Chemical; Cox et al., 2013) was given before the session. For predator odor, 2,3,5-trimethyl-3-thiazoline (TMT; 30 μL; 1% v/v ddH2O) was placed in front each head-restrained mouse on a gauze pad inside a container connected to a vacuum (to control odor spread) for 15 min and removed immediately before testing (King and Becker, 2019). TMT, a synthetically derived component of fox feces, was chosen to test for stress-triggering effects given its ethological relevance and validity (Janitzky et al., 2015). For saline-primed reinstatement, 0.9% NaCl (10 mL/kg, i.p.) was delivered immediately before the session. Animals were re-extinguished to criteria between reinstatement tests. Due to initial piloting, not all mice that completed acquisition and extinction underwent each of the 5 reinstatement tests.

#### Cocaine self-administration

Animals underwent 14 days of 2 hr cocaine (1 mg/kg/12.5 μl cocaine infusion; 40 infusion maximum) self-administration sessions on an FR1 schedule of reinforcement (similar to that described in freely moving mice; Heinsbroek et al., 2017). The criteria for extinction were defined in the same manner as for heroin (see above), and no mice were excluded for failure to reach criteria. Mice underwent reinstatement testing using a counterbalanced design similar to the methods described above for heroin self-administration reinstatement testing, with the exception of drug-primed reinstatement involving cocaine (5 mg/kg, i.p.) rather than heroin injections immediately before the session.

#### Saline self-administration

Mice underwent 14 days of 2 hr saline (12.5 μl infusions; 40 infusion maximum) self-administration sessions on an FR1 schedule of reinforcement. A subset of mice only underwent 13 days of acquisition and did not undergo further testing due to initial piloting (n=4). After completion of acquisition, saline self-administration mice underwent at least 10 days of extinction. Due to low responding during acquisition and thus an inability to “extinguish” lever pressing, mice were immediately tested after the 10^th^ day of extinction. Mice underwent reinstatement testing using a counterbalanced design identical to the methods described above for heroin self-administration experiments.

### Multiphoton imaging

We visualized and longitudinally tracked dmPFC neurons throughout heroin self-administration, extinction, and reinstatement using a multiphoton microscope (Bruker Nano Inc.) equipped with: a hybrid scanning core with galvanometers and fast resonant scanners (30 Hz; we recorded with 4 frame averaging to improve spatial resolution), multi-alkali photo-multiplier tubes and GaAsP-photo-multiplier tube photo detectors with adjustable voltage, gain, and offset features, a single green/red NDD filter cube, a long working distance 20x air objective designed for optical transmission at infrared wavelengths (Olympus, LCPLN20XIR, 0.45NA, 8.3mm WD), a software-controlled modular XY stage loaded on a manual z-deck, and a tunable Insight DeepSee laser (Spectra Physics, laser set to 930nm, ~100fs pulse width). Following imaging sessions, raw data were converted into an hdf5 format for motion correction using SIMA (Kaifosh et al., 2014). Fluorescence traces were extracted from the datasets using Python (Namboodiri et al., 2019; Otis et al., 2019).

### Data collection and statistics

Parameters for behavioral sessions were set using a custom MATLAB graphical user interface that controlled an Arduino and associated electronics. Data were recorded and extracted using MATLAB, analyzed and graphed using GraphPad PRISM (version 8), and illustrated using Adobe Illustrator. Analysis of variance (ANOVA; 2-way or 3-way) or t-tests (paired) were used to analyze data collected for the heroin, cocaine, and saline self-administration experiments. Independent variables included lever (active vs. inactive), day (e.g., extinction vs. reinstatement), and sex (male vs. female). Additionally, a ‘lever discrimination index’ was determined by calculating the ratio of active versus inactive lever presses contributing to total lever presses. The lever discrimination index could not be calculated for saline self-administration experiments as not all mice pressed levers during later acquisition tests. Sidak’s post-hoc tests were used following significant main effects and interactions within the ANOVA analyses.

## RESULTS

### Head-restrained heroin self-administration

Following recovery from surgery and habituation, male and female mice (n=28; 13 males and 15 females) began head-restrained heroin self-administration (Figure 1A-B). Mice learned to reliably press the active lever more than the inactive lever across acquisition (Figure 1C). A mixed-model two-way ANOVA revealed a day by lever interaction (F(13,694)=10.25, *p*<0.001), and post-hoc tests revealed that mice pressed the active lever more than the inactive lever during mid and late (days 5-14; *p*s<0.001) but not early (day 1-4; *ps*>0.05) acquisition sessions. During acquisition, male and female mice did not differ in lever pressing (**Supplemental Figure 1A**), as a three-way ANOVA revealed that there were no effects of sex (F-values<1.5; *p*s>0.1). Heroin self-administration mice were able to discriminate between the active and inactive lever starting from the first day of acquisition (Figure 1D). A mixed-model two-way ANOVA revealed a day by lever interaction for the lever discrimination score (F(13,694)=5.675; *p*<0.001), and post-hoc tests showed that mice significantly discriminated between the active and inactive levers on all days of acquisition (*p*s<0.001). The lever pressing behavior resulted in mice reaching the maximum number of heroin infusions during all sessions (Figure 1E), with male and female heroin mice receiving a similar number of infusions (**Supplemental Figure 1B**) throughout acquisition (two-way ANOVA, effects of sex: F-values<1.0, *p*s>0.5). Thus, mice rapidly and readily acquire heroin self-administration while head restrained.

**Figure 1.**
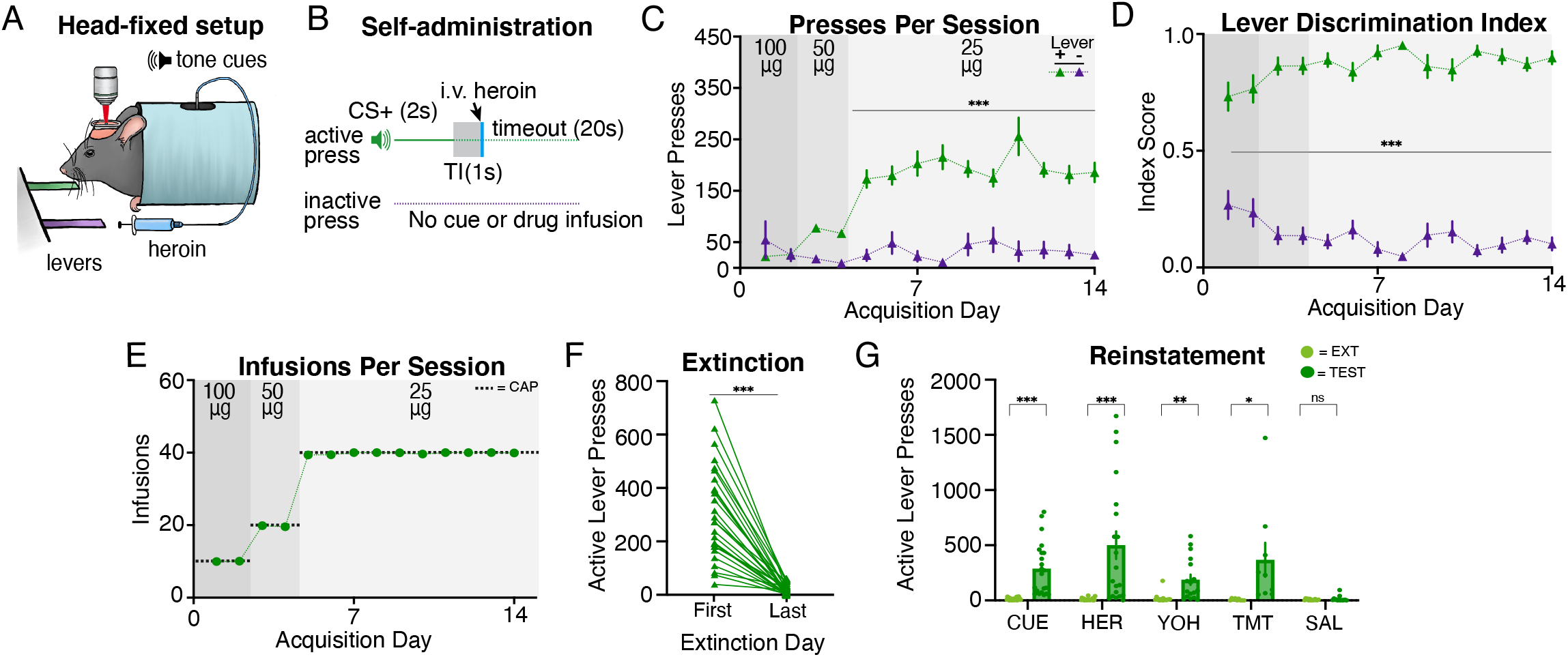
Head-restrained mice self-administer heroin and display reinstatement of heroin seeking. (**A**) Illustration of head fixation for heroin self-administration experiments. Active lever presses resulted in a tone cue that predicted intravenous heroin at doses of 100μg/kg (day 1-2), 50μg/kg (day 3-4) or 25μg/kg (day 5-14). (**B**) Schematic for heroin self-administration experiments, where active but not inactive lever presses resulted in a tone cue (2s), a gap in time (TI, trace interval; 1s), an infusion of heroin, and a timeout period (20s). (**C**) Mice pressed the active lever significantly more than the inactive lever, starting on day 5 of acquisition. (**D**) Mice discriminated between active and inactive levers during all days of acquisition. (**E**) Grouped data showing that mice self-administered the maximum number (CAP) of heroin infusions during each acquisition session, resulting in 1 mg/kg of heroin per session. (**F**) During extinction, mice decreased active lever pressing such that extinction criteria were reached. (**G**) Data showing that mice displayed cue-, heroin-, yohimbine-, and predator odor (TMT)-induced reinstatement of active lever pressing as compared with the previous extinction test. Mice did not reinstate following an injection of saline.

Following acquisition, mice underwent extinction training until criteria were met. A paired t-test confirmed mice significantly decreased active lever pressing from the first day of extinction to the last (Figure 1F; t(27)=8.954; *p*<0.001), with male and female mice displaying equivalent active lever presses on each day (**Supplemental Figure 1C**; two-way ANOVA, effects of sex: F-values<0.3, *p*s>0.5). Next, mice underwent cue- (n=22; male=10, female=12), heroin- (n=20; male=8, female=12), yohimbine- (n=17; male=8, female=9), predator odor- (n=9; male=4, female=5), and/or saline- (n=14; male=7, female=7) induced reinstatement tests (Figure 1G). Paired t-tests revealed that mice significantly increased active lever pressing during cue- (t(21)=5.407; *p*<0.001), heroin- (t(19)=3.966; *p*<0.001), yohimbine- (t(16)=3.866; *p*=0.001), and predator odor-induced (t(9)=2.333; *p*=0.048) reinstatement tests relative to responding during the most recent extinction session. However, the mice did not exhibit reinstatement following an injection of saline (t(13)=0.889; *p*=0.389). We did not observe sex differences in reinstatement responding (**Supplemental Figure 1H**; two-way ANOVAs, effects of sex: F-values<2.4, *p*s>0.1). Overall, we find that head-restrained animals readily self-administer heroin, extinguished lever pressing, and display reinstatement of heroin seeking similar to freely moving mice (Corre et al., 2018).

### Head-restrained cocaine self-administration

Following surgery and habituation, mice (n=8; 4 males and 4 females) underwent head-restrained cocaine self-administration (Figure 2A-B). Mice did not press the active lever significantly more than the inactive lever during acquisition, likely due to variability between animals (Figure 2C). A mixed-model two-way ANOVA revealed that there was only a trend for day by lever interaction (F(13,182)=1.743, *p*=0.056) and post-hoc tests demonstrated there were no significant differences between active and inactive lever pressing throughout acquisition (days 1-14, *ps*>0.05). Additionally, male and female mice were found to press active and inactive levers similarly throughout acquisition (**Supplemental Figure 1E**; three-way ANOVA, effects of sex: F-values<0.5, *p*s>0.8). However, the lever discrimination index revealed that mice were able to significantly discriminate between inactive and active levers (Figure 2D) later in acquisition. A two-way ANOVA of lever discrimination revealed a trend for day by lever interaction (F(13,182)=1.750, *p*=0.054), and post-hoc tests revealed that cocaine mice significantly discriminated between levers during late acquisition (days 9, 11-14; *ps*<0.05), but not early in acquisition (days 1-8, 10; *ps*>0.05). Importantly, lever-pressing behavior resulted in mice reaching the maximum amount of cocaine infusions during each session (Figure 2E), with male and female mice receiving a similar number of infusions (**Supplemental Figure 1F**) during acquisition (two-way ANOVA, effects of sex: F-values<1.3, *p*s>0.3). Thus, mice reliably acquire cocaine self-administration behavior while head restrained.

**Figure 2.**
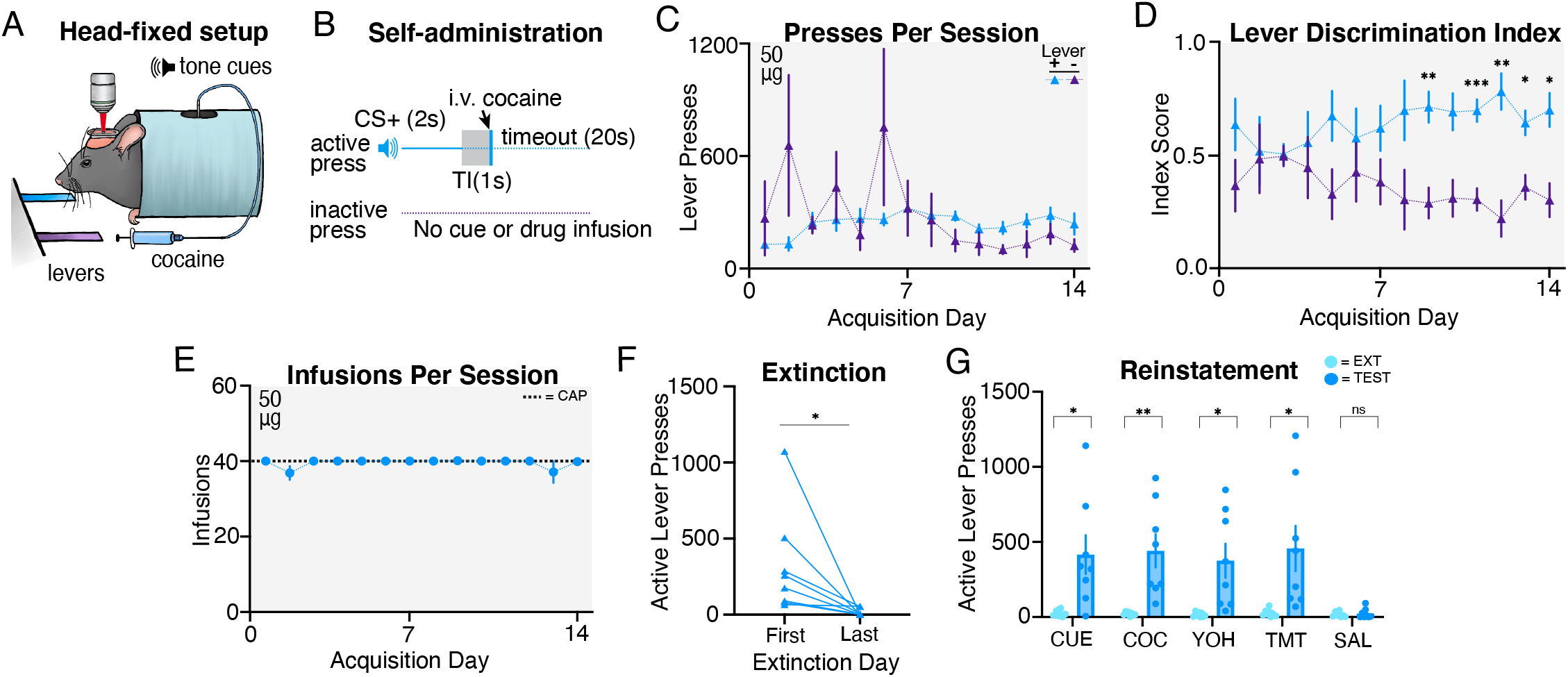
Head-restrained mice self-administer cocaine and display reinstatement of cocaine seeking. (**A**) Illustration of head fixation for cocaine self-administration experiments. Active lever presses resulted in a tone cue that predicted intravenous cocaine (50μg/kg on all days). (**B**) Schematic for cocaine self-administration experiments, where active but not inactive lever presses resulted in a tone cue (2s), a gap in time (TI, trace interval; 1s), an infusion of cocaine, and a timeout period (20s). (**C**) Group data suggest that there was a trend for more pressing for the active lever versus the inactive lever during later days of acquisition (day 9-14). (**D**) Mice significantly discriminated between active and inactive levers during the later days of acquisition (day 9-14). (**E**) Grouped data showing that mice generally self-administered the maximum number (CAP) of cocaine infusions during each acquisition session. (**F**) During extinction, all mice decreased active lever pressing and reached extinction criteria. (**G**) Mice displayed cue-, cocaine-, yohimbine-, and predator odor (TMT)-induced reinstatement of active lever pressing as compared with the previous extinction test. Mice did not reinstate following an injection of saline.

Following cocaine self-administration, mice underwent extinction until criteria were met (n=8; 4 males and 4 females). A paired t-test confirmed that mice significantly decreased active lever pressing from the first day of extinction to the last day (Figure 2F; t(7)=82.461; *p*=0.043), with males and females displaying a similar number of active lever presses on each day (**Supplemental Figure 1G**; two-way ANOVA, effects of sex: F-values<0.4, *p*s>0.5). Next, mice underwent cue-, cocaine-, yohimbine-, predator odor-, and saline-induced reinstatement tests (Figure 2G). Paired t-tests revealed that, compared to the respective prior day’s extinction responding, mice significantly increased active lever pressing during cue- (t(7)=2.954; *p*=0.021), cocaine- (t(7)=3.638; *p*=0.008), yohimbine- (t(7)=3.240; *p*=0.014), and predator odor-induced (t(7)=2.848; *p*=0.025) reinstatement tests. However, mice did not exhibit reinstatement following an injection of saline (t(7)=0.455; *p*=0.663). We did not observe sex differences during the reinstatement tests (**Supplemental Figure 1H**; two-way ANOVAs, effects of sex: F-values<0.9, *p*s>0.3), although there was a trend between males and females following the control injection of saline (effect of sex: F(1,6)=5.644, *p*=0.055; sex by day interaction: F(1,6)=2.979, *p*=0.135) reinstatement tests. Overall, we find that head-restrained animals self-administer cocaine, extinguished lever pressing, and display reinstatement of cocaine seeking similar to freely moving mice (Heinsbroek et al., 2017).

### Head-restrained saline self-administration

To confirm that head-restrained drug self-administration mice are learning to press the active lever for the drug rewards, and not for the tone CS alone, we included a head-restrained saline self-administration control group (n=8; 5 males and 3 females). Following recovery and habituation, mice began head-restrained saline self-administration (Figure 3A-B), using a similar behavioral protocol as the previous heroin and cocaine self-administration experiments. Saline self-administration mice did not press the active lever more than the inactive (Figure 3C), as a two-way ANOVA confirmed there was no effect of lever (F(1,14)=0.001, *p*=0.096). Furthermore, although there was a day by lever interaction (F(13,168)=2.697, *p*=0.002). post-hoc analyses revealed no significant differences in active versus inactive lever pressing on any session. Unlike the heroin or cocaine self-administration mice, saline self-administration mice did not reach the maximum number of saline infusions throughout acquisition (Figure 3D). Following acquisition, a subset of saline self-administration mice (n=4) underwent ten days of extinction before reinstatement testing (Figure 3E). Paired t-tests revealed that saline self-administration mice did not increase active lever pressing during cue- (t(3)=0.858; *p*=0.454), heroin- (t(3)=1.894; *p*=0.155), yohimbine- (t(3)=2.294; *p*=0.106), or saline-induced reinstatement tests (t(3)=1.127; *p*=0.342). Overall, we find that head-restrained mice readily self-administer drugs of abuse but not saline.

**Figure 3.**
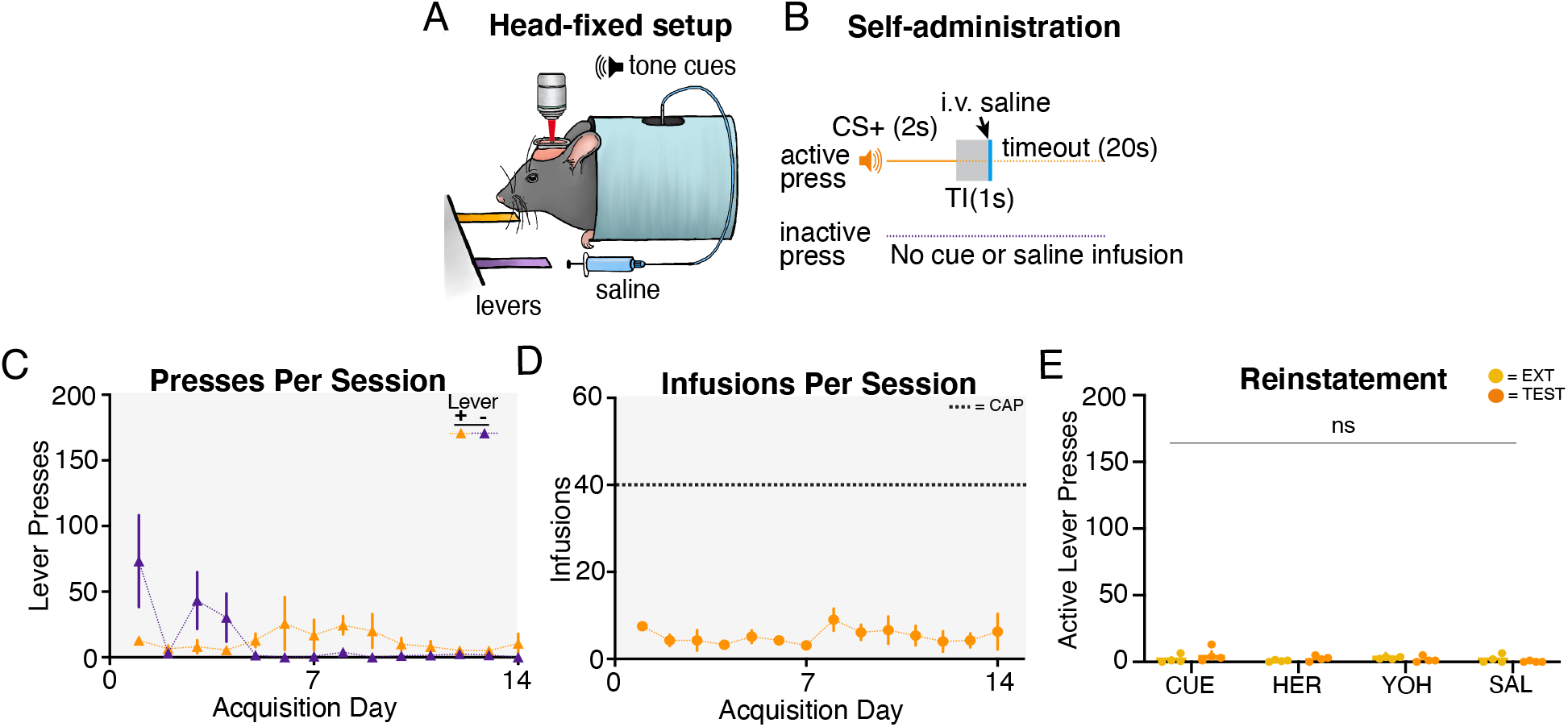
Head-restrained mice do not self-administer saline or display saline seeking. (**A**) Illustration of head fixation for saline self-administration experiments. (**B**) Schematic for saline self-administration experiments, where active but not inactive lever presses resulted in a tone cue (2s), a gap in time (TI, trace interval; 1s), an infusion of saline, and a timeout period (20s). (**C**) Group data showing that mice did not press the active lever more than the inactive lever during acquisition. (**D**) Grouped data showing that mice did not self-administer the maximum number (CAP) of saline infusions during acquisition. (**E**) Mice did not increase lever pressing during cue-, heroin-, yohimbine-, predator odor (TMT)-, or saline-induced reinstatement tests as compared with the previous extinction test.

### Visualization of dmPFC excitatory neurons throughout heroin self-administration

We next evaluated the feasibility of combining head-restrained drug self-administration with multiphoton imaging. Using a virus encoding the calcium indicator GCaMP6s (Chen et al., 2013) in combination with two-photon microscopy, we visualized the calcium dynamics of single dmPFC excitatory output neurons from a mouse *in vivo* (Figure 4A-B). The activity of individual neurons could be resolved and longitudinally tracked throughout each drug self-administration, extinction, and reinstatement session. Standard deviation projection images from time series data reveal the fluorescence the tracked cells during each phase of acquisition (early, middle, late), extinction (early, late), and cue-, drug-, predator odor-, yohimbine-, and saline-induced reinstatement tests (Figure 4C-L). Overall, we are able to simultaneously and longitudinally visualize the activity of single, genetically-defined neurons from the onset of drug use to reinstatement using the head-restrained drug self-administration approach.

**Figure 4.**
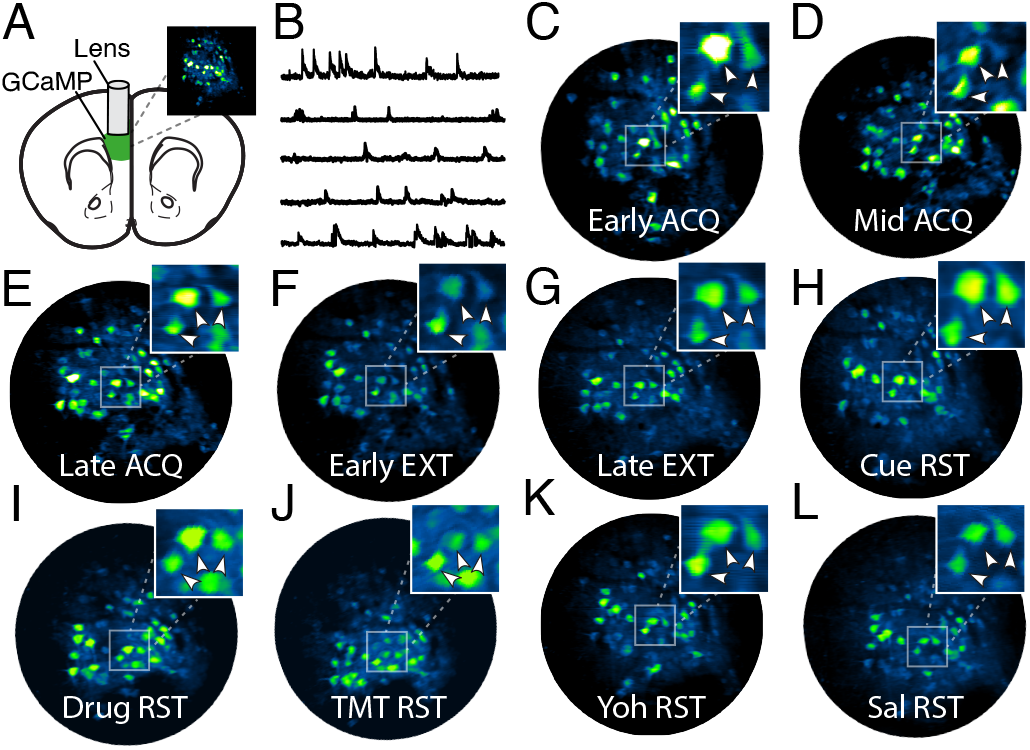
Single cell tracking from the onset of heroin use to reinstatement. (**A**) Surgical strategy and (**B**) example *in vivo* fluorescence dynamics among GCaMP6s-expressing dmPFC excitatory neurons. (**C-L**) Standard deviation projection images of dmPFC neurons reveal that a subset of neurons can be tracked throughout acquisition (C-E), extinction (F-G), and reinstatement sessions (H-L). Arrowheads point to tracked neurons. ACQ, acquisition; EXT, extinction; RST, reinstatement.

## DISCUSSION

Here we demonstrate that head-restrained mice will readily self-administer drugs of abuse. We find this to be the case regardless of drug class (i.e., opioids and psychostimulants) and that seeking behavior is specific to rewarding stimuli (i.e., not saline). Furthermore, we demonstrate that mice will extinguish drug seeking if lever pressing is no longer reinforced, whereas mice will resume seeking upon re-exposure to stimuli known to provoke relapse in humans (i.e., drug-associated cues, the drug itself, and stressors; Kalivas and Volkow, 2005). These findings indicate that our head-restrained procedure shares similar construct and predictive validity as drug self-administration assays in freely moving rodents (Epstein et al., 2006; SanchisLJSegura and Spanagel, 2006). Furthermore, this study demonstrates the feasibility of of combining preclinical drug self-administration experiments with novel technologies, such as multiphoton microscopy, that require head immobilization.

### Tracking activity in cell-type specific neurons from the onset of drug use to relapse

The development of SUD involves complex and long-lasting changes to activity in brain reward circuitry (Koob and Volkow, 2016), but how these adaptations develop in cell-type specific neurons and predict relapse vulnerability is unknown. With the advent of multiphoton microscopy (Denk et al., 1990) along with genetically encoded calcium indicators (Chen et al., 2013), we can now longitudinally measure activity in deep brain, cell-type specific neurons for weeks to months in awake, behaving animals (Namboodiri et al., 2019; Otis et al., 2017, 2019; Rossi et al., 2019). This combinatorial approach can be exploited to monitor activity not only in hundreds to millions of neuronal cell bodies simultaneously (Dombeck et al., 2007; Kim et al., 2016), but also in dendrites (Lavzin et al., 2012), dendritic spines (Chen et al., 2011), and axons (Lovett-Barron et al., 2014; Otis et al., 2019). Although calcium indicators are by far the most commonly used for visualizing activity, other sensors are available and under continued development for visualization of ground-truth voltage (Gong et al., 2015), neurotransmitter release and binding (Jing et al., 2018; Marvin et al., 2018, 2019; Nguyen et al., 2010); molecular signaling (Greenwald et al., 2018; Muntean et al., 2018), and more (Aper et al., 2016; Díaz-García et al., 2019; Marvin et al., 2011). These powerful technologies could be combined with drug self-administration studies in head-restrained mice to provide unprecedented insight into the abnormal neural circuit activity patterns that emerge from the onset of drug use to relapse.

### Tracking morphological plasticity in cell-type specific neurons from the onset of drug use to relapse

Structural plasticity is a critical mechanism that contributes to dysfunctional neural circuit activity patterns in SUD. This plasticity is apparent at the level of dendrites and dendritic spines, axon terminals, astrocytes, extracellular matrices, and more (for review, see Kruyer et al., 2020; Russo et al., 2010; Spiga et al., 2014; Wolf, 2016). Although these structural modifications are rapidly and dynamically regulated throughout drug use, withdrawal, abstinence, and relapse – scientists have been limited to *ex vivo* histological analysis and between subjects comparisons to study these adaptations. Multiphoton imaging, in combination with genetically encoded fluorophores for visualization of cell morphology, allows measurement of structural plasticity in precisely defined cell types in awake, behaving animals (Moda-Sava et al., 2019; Muñoz-Cuevas et al., 2013). Thus, by combining multiphoton microscopy with head-restrained drug self-administration experiments, we are no longer limited to snapshots of the addicted brain. Rather, we can perform longitudinal recordings to investigate the morphological adaptations that arise in precisely defined neural circuit elements from the onset of drug use to relapse.

### Evaluating the function of neuronal ensemble activity patterns and morphological plasticity in drug use and seeking

Studies using multiunit recordings and calcium imaging technologies often reveal complex activity patterns within single brain regions during natural reward (Otis et al., 2017, 2019) and drug reward-seeking (Ahmad et al., 2017; Drouin and Waterhouse, 2004; Siciliano et al., 2019). These heterogeneous activity patterns are not just common at the population level, but also at the level of neuronal subpopulations, as recordings from genetically defined or projection-specific neurons also reveal profound cell-to-cell variability (Al-Hasani et al., 2015; Dembrow and Johnston, 2014; Seong and Carter, 2012; Yang et al., 2018). The inherent complexity of neuronal circuit activity patterns has made it difficult to selectively manipulate unique neuronal ensembles to determine their function for behavior. However, existing and emerging technologies that combine multiphoton microscopy with optogenetics now allow for manipulation of activity in experimenter-defined neurons in 3-dimensional space (Marshel et al., 2019; Yang et al., 2018). Furthermore, multiphoton laser-directed lesion experiments have been employed at the level of cell bodies, axons, dendrites, and dendritic spines *in vivo* (Allegra Mascaro et al., 2010; Canty et al., 2013; Hill et al., 2017; Park et al., 2019). These technologies could now be combined with drug self-administration studies to identify the function of unique neuronal ensembles and morphological plasticity for drug use and seeking.

### Limitations and future directions

A limitation of our protocol is that head restraint is likely to be stress provoking. Chronic stress can lead to escalation of drug intake (Mantsch and Katz, 2007), which may produce behavioral phenotypes akin to those observed in animals with a greater history of intake (i.e., “long-access” or “extended-access”, rather than “short-access”, animals; Mantsch et al., 2016). Future modifications to our approach could be beneficial for reducing the possible effects of stress on drug seeking and could allow for improved behavioral resolution within the task itself. For example, inclusion of a running wheel, treadmill, or trackball would provide an avenue for mice to move while head restrained and would also provide a behavioral readout of locomotor activity – a behavioral variable that is often studied due to its robust modulation by repeated drug use and correlation with addiction vulnerability (Piazza et al., 1989; Zhou et al., 2019). Altogether, the influence of restraint-related stress should be acknowledged and considered when designing drug self-administration experiments in head-restrained animals. Moreover, future adaptations of the assay could improve its strength for studying the neural circuit underpinnings of SUD.

### Concluding remarks

By coupling multiphoton imaging with simultaneous intravenous drug self-administration, we can now characterize and manipulate adaptations in neuronal circuits that evolve from the onset of drug use to relapse. The replication of effects observed in freely moving animals indicates that our head-restrained approach could build upon, rather than diverge from, the foundation provided by the years of research conducted with the drug self-administration paradigm. Most importantly, this could accelerate discovery of novel therapeutic interventions for SUD.

## CONTRIBUTIONS

KMV, EMD, and JMO designed the experiments and wrote the manuscript. KMV, EMD, RIG, KTW, EVR, CWB, PNS, and ITP performed the experiments. ACB and PWK provided intellectual prsupport and training for self-administration studies.

## ACKNOWLEDGMENTS

The authors would like to thank Dr. Madalyn Hafenbreidel, Dr. Carmela Reichel, Dr. Jakie McGinty, Dr. Howard Becker, Eric Dereschewitz, and Maribel Vasquez-Silva for technical advice and support.

---

**Supplemental Figure 1. Sex comparison across head-fixed drug self-administration experiments.**

*Heroin self-administration.* (**A**) Male and female mice did not differ in active and inactive lever pressing during acquisition of heroin self-administration. (**B**) Male and female mice did not differ in the number of i.v. heroin infusions received during acquisition. (**C**) Male and female mice did differ in active lever pressing rates during extinction of heroin seeking. (**D**) Male and female mice did not display differences during cue-, heroin-, yohimbine-, or predator odor (TMT)-, or saline-induced reinstatement tests.

*Cocaine self-administration.* (**E**) Male and female mice did not differ in active and inactive lever pressing during acquisition of cocaine self-administration. (**F**) Male and female mice did not differ in the number of i.v. cocaine infusions received during acquisition. (**G**) Male and female mice did differ in active lever pressing rates during extinction of cocaine seeking. (**H**) Male and female mice did not display differences during cue-, cocaine-, yohimbine-, or predator odor (TMT)-, or saline-induced reinstatement tests.

