## Supplementary figures and images for "Drug self-administration in head-restrained mice for simultaneous multiphoton imaging"

### Supplemental Figure 1A

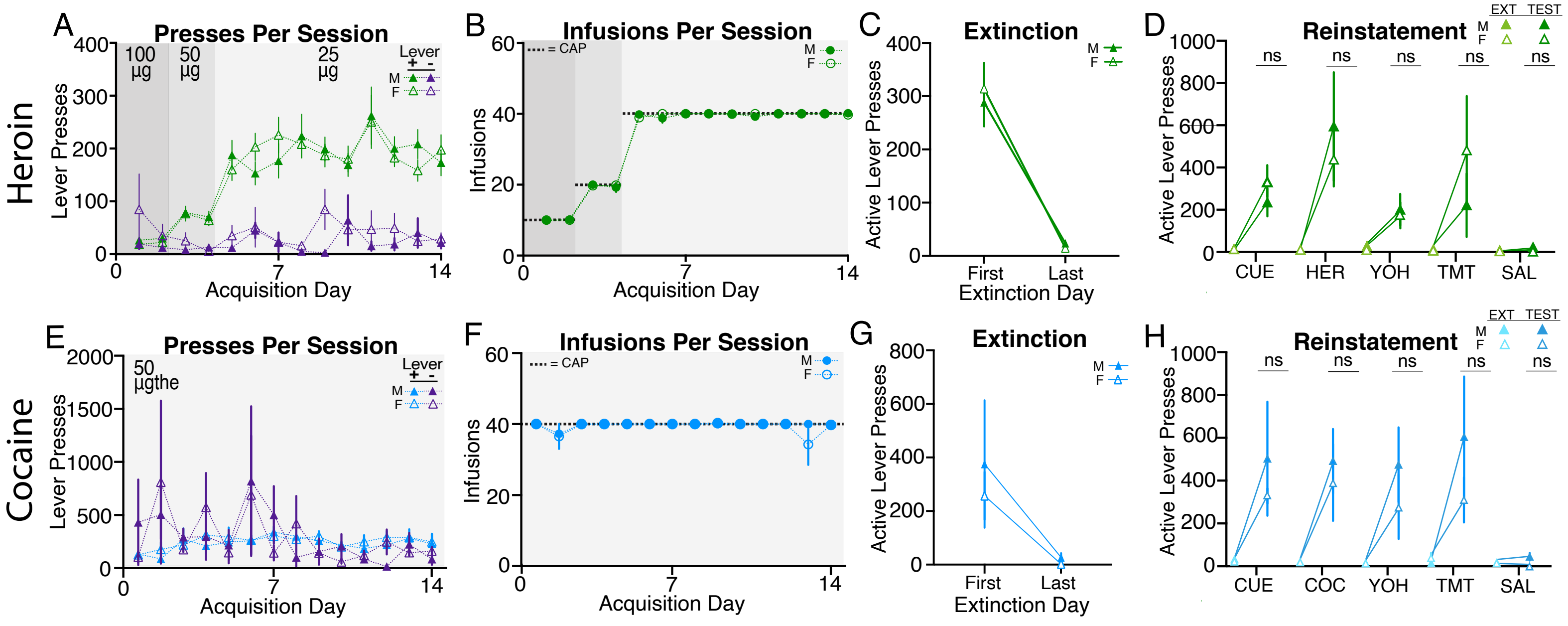
